# A theory of the use of information by enemies in the predator-prey space race

**DOI:** 10.1101/2020.01.25.919324

**Authors:** Rémi Patin, Daniel Fortin, Simon Chamaillé-Jammes

## Abstract

We currently lack a comprehensive theory about how behaviourally responsive predators and prey use the information they acquire about the environment and each other’s presence while engaged in the ‘space race’. This limits our understanding of the role of behaviour in trophic relationships and our ability to predict predators and prey distributions. Here we combined a simulation model with a genetic algorithm to discover how predators and prey behaving optimally should use information in environments with different levels of heterogeneity in prey forage distribution and prey vulnerability. Our results demonstrate the key role of movement unpredictability in successful strategies for both predators and prey, supporting the ‘shell-game’ hypothesis. We however reveal striking differences between predators and prey in the magnitude of this unpredictability, and in how it varies with the environment. Our work offers a robust theoretical framework to better understand predator-prey space use and interpret empirical studies.

## Introduction

Predators and prey can be viewed as engaged in a ‘space race’ (Sih 1984). Prey select patches offering a favourable balance between foraging opportunities and predation risk (Gilliam & Fraser 1987; Brown & Kotler 2004), while predators search for patches where they have the greatest chance of capturing prey (Kacelnik et al. 1992; Hopcraft et al. 2005). Not only prey and predators are most often on the move, but prey and predators generally live in environments where resources are heterogeneously distributed as they get locally depleted, and renew. To succeed in such dynamic environments, both prey and predators should have evolved cognitive abilities to perceive, memorize and later use information about the environment and the distribution of their enemies in the space race.

Simple models have shown that the use of past information can greatly improve foraging success of individuals in heterogeneous environments by reducing their search time and improving the timing of their visits to patches, even when competition decreases resource predictability (Riotte-Lambert et al. 2015, 2017)(see also (Merkle et al. 2014) for an empirical demonstration). Selection pressure for the evolution of mnesic capacity is likely strong and, accordingly, memory-driven space use is increasingly demonstrated (Fagan et al. 2013; Merkle et al. 2019). The use of memory could however make predators and prey more predictable, allowing prey to develop spatial avoidance strategies, and predators to search more efficiently. For instance, a prey foraging on a limited set of rich resource patches and traveling between them in virtually the same order (‘traplining’, Riotte-Lambert et al. 2016) may become highly predictable by its predator. Even if prey move less deterministically, only avoiding patches with little resources, predators could perceive the intrinsic quality of the patches for the prey, memorize the location of food-rich patches, and focus their search among them (‘leap-frogging’, Sih 1998). Meanwhile, prey could also learn about this now predictable distribution of predators, and start avoiding the resource-rich, but risky, patches. Predators would thus be unlikely to persist with such a strategy. Generally, predators and prey face the dilemma that, while the use of information could apparently be beneficial to them, its use could also make them more predictable, thereby providing some benefits to their enemies in the predator-prey space race.

Under these conditions, the predator-prey race could rather be viewed as a ‘shell game’ in which “predators search for elusive prey, and prey stay on the move to remain elusive” (Mitchell & Lima 2002). Such view emphasizes that randomness could be an integral part of optimal predator and prey movement strategies (Mitchell & Lima 2002; Mitchell 2009). Randomness may, however, loosen the link between an individual and its resources, reducing the benefits of memory use. Mitchell & Lima (2002) and Mitchell (2009)’s models however showed that it can ultimately be beneficial for prey to move randomly, and thus be unpredictable, when the predator can memorize the locations of past encounters. In such situation, the predator would be able to concentrate its search on prey-rich patches if those prey only focused on forage-rich patches and move predictably. In these models, however, the prey had no memory, and their movement strategies were compared only across various types of random movement. Using a different model, Bracis et al. (2018) showed that prey would benefit from using memory under predation risk, especially when predation risk is decoupled from resource distribution as prey can thus avoid predators while foraging in resource-rich areas. In this model, predators could however not adjust to prey distribution. As for prey, it has been argued that predators would gain from being unpredictable (Roth & Lima 2007; Valeix et al. 2011), although this would involve them searching areas where prey are rarer. We are not aware of models predicting the conditions under which randomness in predator movements would benefit them. Empirical demonstrations of the shell game are only emerging (Simon et al. 2019)(but see Laundré 2010), and the optimal level of randomness that prey and predators should optimally use remains unknown. No theory integrating the co-evolution of prey and predator movement strategies has been proposed yet, and we argue this is a fundamental gap in our ability to understand and predict the movement of animals.

Here, we develop theoretical expectations for the optimal use, by behaviourally responsive predators and prey, of randomness and current and past information about prey forage availability and encounters. We study whether these expectations vary with the abundance and spatial heterogeneity of prey forage resources and prey vulnerability across patches.

Overall, we address 3 general questions, for both predators and prey:

1. How important is the information about prey forage availability compared to the information about encounters with enemies?
2. How important is recent vs. older information on both forage availability and encounters with enemies?
3. Is it beneficial to introduce some level of randomness in the movement strategy, and if so how much is optimal?

## Methods

We combined a simulation model and a genetic algorithm to find the optimal movement strategies of both predators and prey. The implementation was done in R (R Core Team 2018) using the Rcpp package (Eddelbuettel & Balamuta 2018).

### Model outline

The model simulates the movement of a fixed number of predators and prey between a set of patches. Patches are characterized by (i) the current amount of forage available to the prey (hereafter referred to as ‘forage’), which increases logistically over time but decreases with prey consumption, (ii) the maximum amount of forage that can exist in the patch and (iii) the vulnerability (i.e. the likelihood of dying) of prey when attacked by a predator or, equivalently, the success rate of attacks of the predators. At each time step, prey forage and accumulate resources. Predators have a probability of encountering a prey that increases with prey density in the patch, but attack only once per time step. Both prey and predators assign weights to each patch: weights represent the assessment of patch quality from past experience, and the weight of a patch is therefore re-evaluated following each visit. Both predators and prey have movement strategies, defined by the values of 4 parameters (α, μ, β, and δ, see ‘Model formulation’ section below), determining whether forage or encounters, and recent or old information for each, are contributing more to the weighting of the patches, and finally whether weights contribute a lot or little to the actual choice of the patch used at the next time step.

### Model Formulation

Central to the model is the fact that each predator and prey store estimates of the quality of patches as weights. For any given individual, the current weight given to patch *i* is:

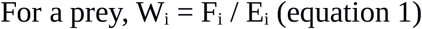

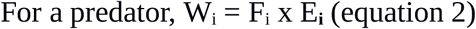

where F_i_ is a state variable indexing the perceived quality of patch *i* with regard to prey forage, and E_**i**_ a state variable indexing the perceived chance of being predated upon (for a prey) or of successfully capturing a prey (for a predator) in patch *i* (more details below).

The model consists in a sequence of time steps, themselves decomposed into consecutive substeps. Parameter values and initialization values of state variables are presented in Supporting Information (SI) 1.

### Substep 1: Calculating forage growth and consumption

This substep takes place for all patches at each time step *t*. The amount of forage levels in patches follows a logistic growth reduced by prey consumption, such as:

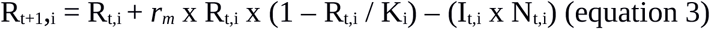

with R_t+1,i_ and R_t,i_ the amount of forage in patch *i* at time *t+1* and *t* respectively, *r_m_* the maximum forage growth rate (constant among patches), K_i_ a forage carrying capacity that can vary among patches, N_t,i_ the number of prey in patch *i* at time *t*, and I_t,i_ the forage intake of a prey in patch *i* at time *t*. Prey forage intake follows a type II functional response, providing there is enough forage in the patch to feed all prey present. If not, forage is equitably shared among all prey present in the patch:

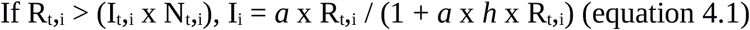

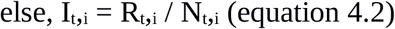

with *a* the attack rate on forage and *h* the handling time.

We assumed that prey consumption could not lead a patch to a stage where it would not regrow, and allowed R_t+1_,i = R_min_ (with R_min_ small, Table S1), if eq. 3 returned R_t+1,i_ < R_min_.

### Substep 2: Estimating encounters

This substep takes place for each predator present in a patch where prey are also present. Each predator search for prey, and encounters each prey sequentially with a constant probability p_e_, until all prey present in the patch have been tested or an encounter occurs. The predator then stops searching during this time step. A prey can however encounter several predators.

### Substep 3: Updating prey patch weights

For each prey, the weight given to the patch *i* currently occupied, and only this one, is updated. The state variable F_i_ indexing the perceived quality, forage-wise, of patch *i*, is updated by integrating the information about the current intake:

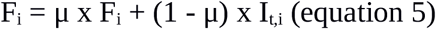

with μ a parameter whose value is within ]0-1[ and determine the relative contribution of recent vs. older information: as the value of μ increases, the relative contribution of old information to F_i_ increases, and F_i_ becomes mostly related to the long-term average intake at the patch, rather than to its recent state.

The state variable E_i_ is updated by integrating the information about predation risk, such as:

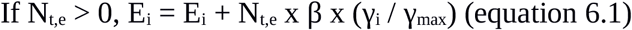

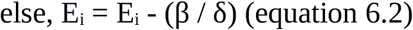

where N_t,e_ is the number of encounters of the prey with predators during this time step, γ_i_ the vulnerability (probability of prey death given an encounter) in patch *i,* and γ_max_ the maximum vulnerability observed among all patches. β represents the importance of information about encounters in determining space use: prey with greater values of β gives lower weights to the patch after an encounter. δ represents the relative contribution of recent vs. older information on encounters in determining E_i_. Specifically, δ x (γ_i_ / γ_max_) time-steps without an encounter in patch *i* are required for the influence of an encounter on the weight of patch *i* to disappear. Therefore, prey with greater δ values give higher importance to old encounters when selecting a patch. Finally, the weight W_i_ is updated following equation 1.

### Substep 4: Updating predator patch weights

For each predator, the weight given to the patch *i* currently occupied, and only this one, is updated. This substep is similar to substep 3 and we only highlight what differs. Firstly, for predators, the state variable indexing prey forage availability F_i_ is updated based on R_t,i_, the amount of forage available in patch *i* at time *t*, rather than on prey forage intake (I_t,i_), which would be harder to evaluate for a predator. Therefore,

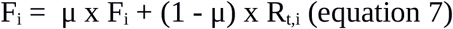

Secondly, as for prey, E_i_ is updated using equations 6.1 or 6.2, but N_t,e_ equals the number of prey encountered by the predator during this time step, which is 0 or 1 as the predator cannot encounter more than one prey during one time step. Finally, the weight W_i_ is updated following equation 2. Note that, in contrast to prey and because of the difference between equations 1 and 2, predators with greater values of β give greater (not lower) weight to the patch after an encounter. β however still index the importance of encounters vs. prey forage in driving patch choice.

### Substep 5: Selecting target patches for predators and prey

This substep is repeated for each predator and each prey. The probability of selecting, at the next time step, patch *i* among patches 1,…, p, is given by:

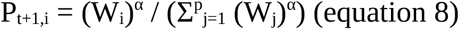

with α a parameter determining to what extent the choice of patches depends on the weights given to patches. If α is very near 0, the individual chooses the next patch virtually at random. If α = 1, the probability for the individual of choosing a specific patch is proportional to the patch weight. If α = 8, the highest value used here, the individual has a very high probability of choosing the patch with the highest weight.

### Fitness calculations

At the end of the simulation, the fitness of each prey is calculated as the product of its total intake and its survival probability. The fitness of each predator is calculated as the number of prey successfully captured.

### Finding optimal strategies

To find the optimal strategies for both predator and prey, we built a genetic algorithm (GA) on top of the model (Hamblin 2013). This approach is especially valuable when brute-force optimization using gradient of parameter values for all parameters across all combinations is not computationally feasible, which was the case for our model. GA relies on a balance between exploration through the generation of new strategies and exploitation through the selection of the best strategies (Hamblin 2013). As such, it can be viewed as mimicking the process of natural selection.

The GA is initialized with a random set of strategies, each strategy being defined by a combination of values given to parameters α, μ, β, and δ. For 4000 generations, we iterate the same operation: (i) evaluation of the fitness of strategies investigated by running the model, (ii) selection and mutation of strategies. First, strategies that will persist are selected. Selection is elitist, so the strategies leading to the 10% highest fitness values are always kept. The other strategies are selected using the k-tournament method, with k = 2 (see Hamblin 2013). After selection of the strategies, each parameter of the strategies has a 5% chance of being subject to mutation, which could be a recombination (2.5%), a small mutation (1.25%) or a large mutation (1.25%). Recombinations replace the current parameter by the one of another strategy taken at random. Small mutations add a small increment to the current parameter value. Large mutations replace the current parameter by a new random value within the parameter range. On completion of the algorithm, a set of strategies remains and an additional series of generations is ran with selection but without mutation, in order to filter the remaining strategies until a single one remains for both predators and prey (Hamblin 2013). Even after a long run-time, a GA may fail to find optimal solutions because of continuous mutations or local optima. In order to check convergence of the algorithm to strategies that would be repeatedly selected in a given environment, we replicated each simulation 3 times. At times, the final value of some parameters differed between replicates. This was the case when strategies with low α values (near-random movement) were efficient, as other parameters therefore did not affect the actual movement of individuals and were subject to drift. Figures in the main text present results for one replicate and those for the 2 others are shown in SI 2.

### Analyses

We compared optimal strategies of predators and prey across environments that differed in (1) the vulnerability in standard patches (4 values); (2) the contrast in vulnerability between standard and riskier patches (5 values); (3) the maximum forage in standard patches (3 values); (4) the contrast in maximum forage between standard and riskier patches (4 values, always >1 so that riskier patches were always intrinsically better than standard patches in terms of prey forage). We tested all combinations of values (see Table S1), yielding a total of 240 different environments. In the main text, we present the distribution of values of strategy parameters as a function of those four environmental characteristics, as well as for environments without predators, to compare prey movement strategy with the one observed under predation risk. In SI 2, we provide a comprehensive report of all results on optimal strategy parameters, as well as on the distribution of predators and prey, for all the environments tested and for the 3 replicates.

## Results

### Movement predictability

In all types of environments, predators are generally much less predictable than prey. Predators generally have α values below 1 (Fig. 1, Fig. S2-1 to S2-5 in SI 2), suggesting significant randomness in their patch choice. Prey movements are much more driven by patch weights (greater α)(Fig. 1, Fig. S2-1 to S2-5). In absence of predators, however, prey would be much more predictable, virtually always choosing the best patches (α ~ 8, Fig. 1, Fig. S2-1 to S2-4).

**Figure 1.**
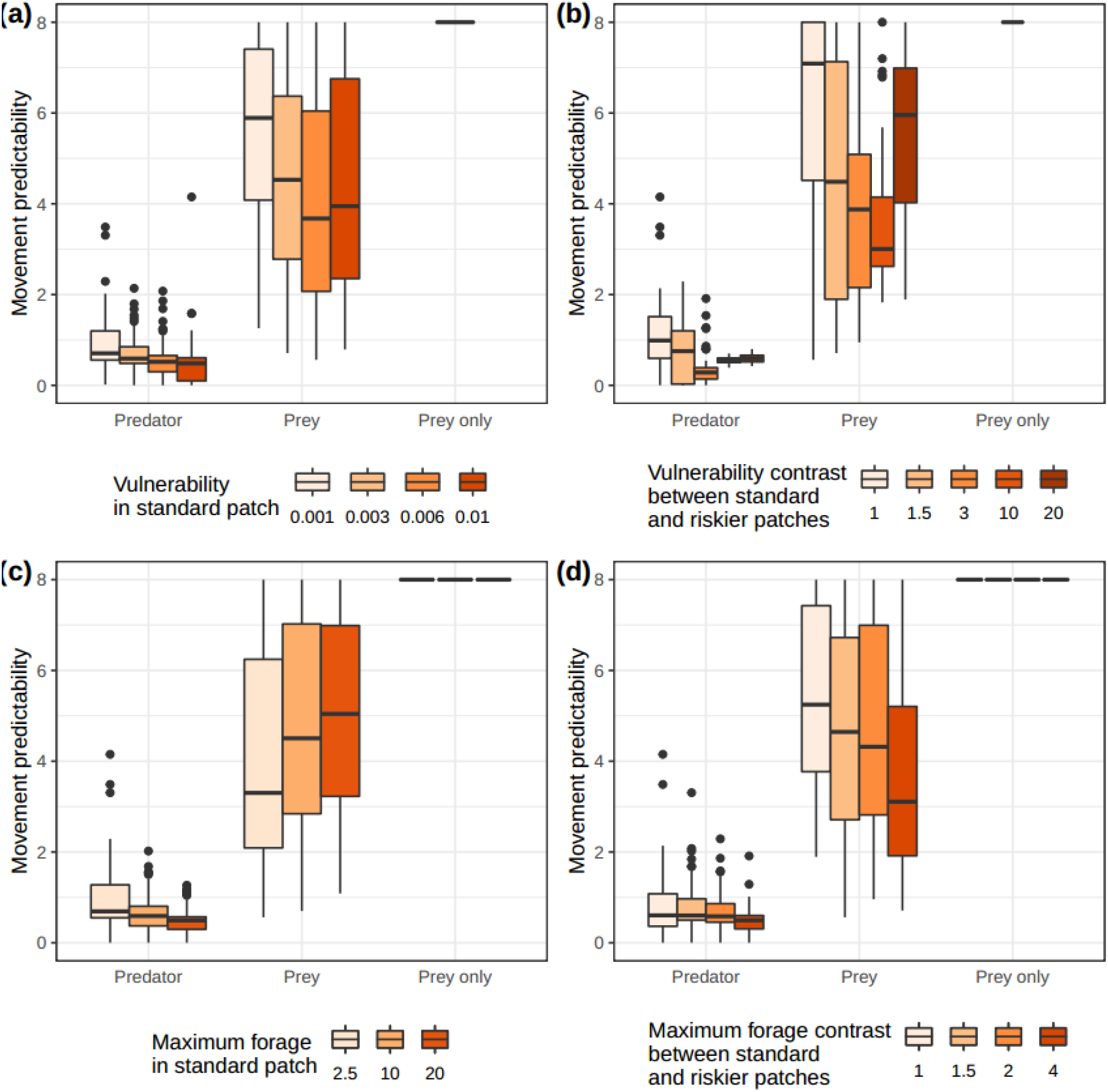
Optimal values, for predators and prey, of parameter α, indexing movement predictability, for varying (a) vulnerability in the standard patches, (b) vulnerability contrast between standard and rich patches, (c) maximum forage in the standard patches and (d) maximum forage contrast between standard and rich patches. Values for environments without predators are also shown (patches were then not assigned vulnerability levels). Lower values correspond to less informed, i.e. more random, movements.

Predator strategies involve greater randomness with higher vulnerability in standard patches (Fig. 1a), higher maximum forage in standard patches (Fig. 1c) and very high forage contrast between standard and riskier patches (Fig. 1d). Predator movement predictability is minimum for intermediate contrast of vulnerability between standard and riskier patches (Fig. 1b). Prey movement strategies involve greater randomness as vulnerability in standard patches increases (Fig. 1a), vulnerability contrast between standard and riskier patches increases (except for highest value, Fig. 1b), maximum forage in standard patches decreases (Fig. 1c), and forage contrast between standard and riskier patches increases (Fig. 1c). In absence of predators, prey were always highly predictable (Fig. 1).

### Relative importance of information about encounters

Keeping in mind that predators use past information less than prey to decide where to move (see above), encounters have a greater importance (greater values of β, Fig. 2, Fig. S2-6 to S2-10 in SI 2) for predators’ assessment of patch quality than for prey. Predators grant very high importance to information about encounters in all environments except when the contrast of prey vulnerability is very low between standard and riskier patches (Fig. 2, Fig. S2-6 to S2-10). Prey generally give little importance to information about encounters, but this information becomes slightly more important as vulnerability in the standard (i.e. safest) patches increase (Fig. 2a, Fig. S2-6), the contrast in vulnerability between standard and riskier patches increases (Fig. 2b, Fig. S2-7), and the contrast in maximum forage between standard and riskier patches increases (Fig. S2-9).

**Figure 2.**
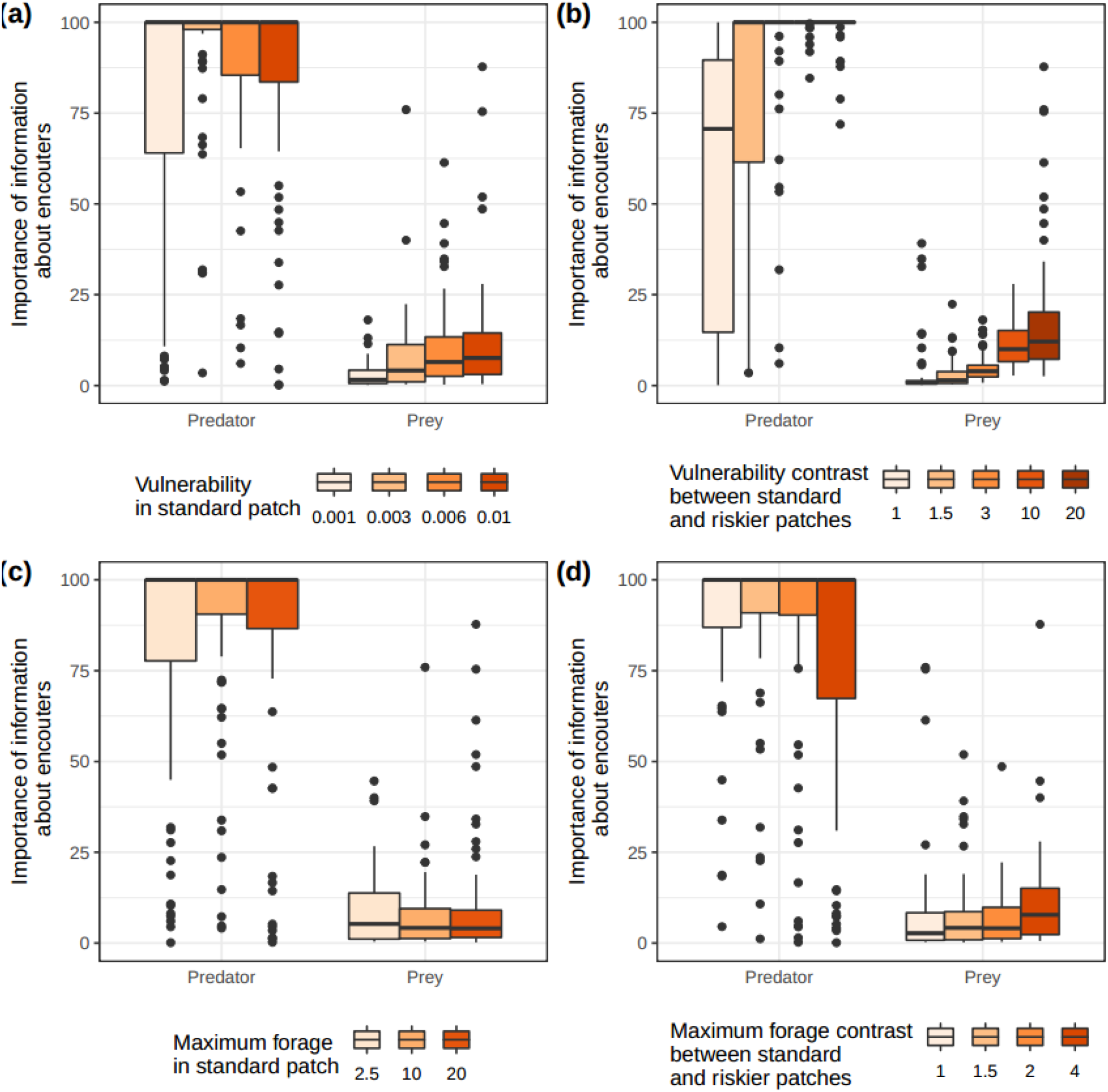
Optimal values, for predators and prey, of parameter β, indexing the relative importance of information about encounters, for varying (a) vulnerability in the standard patches, (b) vulnerability contrast between standard and rich patches, (c) maximum forage in the standard patches and (d) maximum forage contrast between standard and rich patches. Lower values correspond to lower importance given to encounters, relative to prey forage availability.

### Relative importance of recent vs. older information about encounters

Generally, predators and prey differ little in how they integrate information about recent and older encounters when determining patch weights (Fig. 3, Fig. S2-11 to S2-15 in SI 2). Both predators and prey grant greater importance to old encounters as vulnerability in standard patches increases (Fig. 3a, Fig. S2-11), and as the vulnerability contrast between standard and riskier patches increases (Fig. 3b, Fig. S2-12). When this vulnerability contrast becomes very large, prey use information about old encounters more than predators (Fig. 3b, Fig. S2-12). Prey forage availability has a limited effect on prey and predators’ relative use of information about recent and old encounters (Fig. 3c, 3d, Fig. S2-13, S2-14).

**Figure 3.**
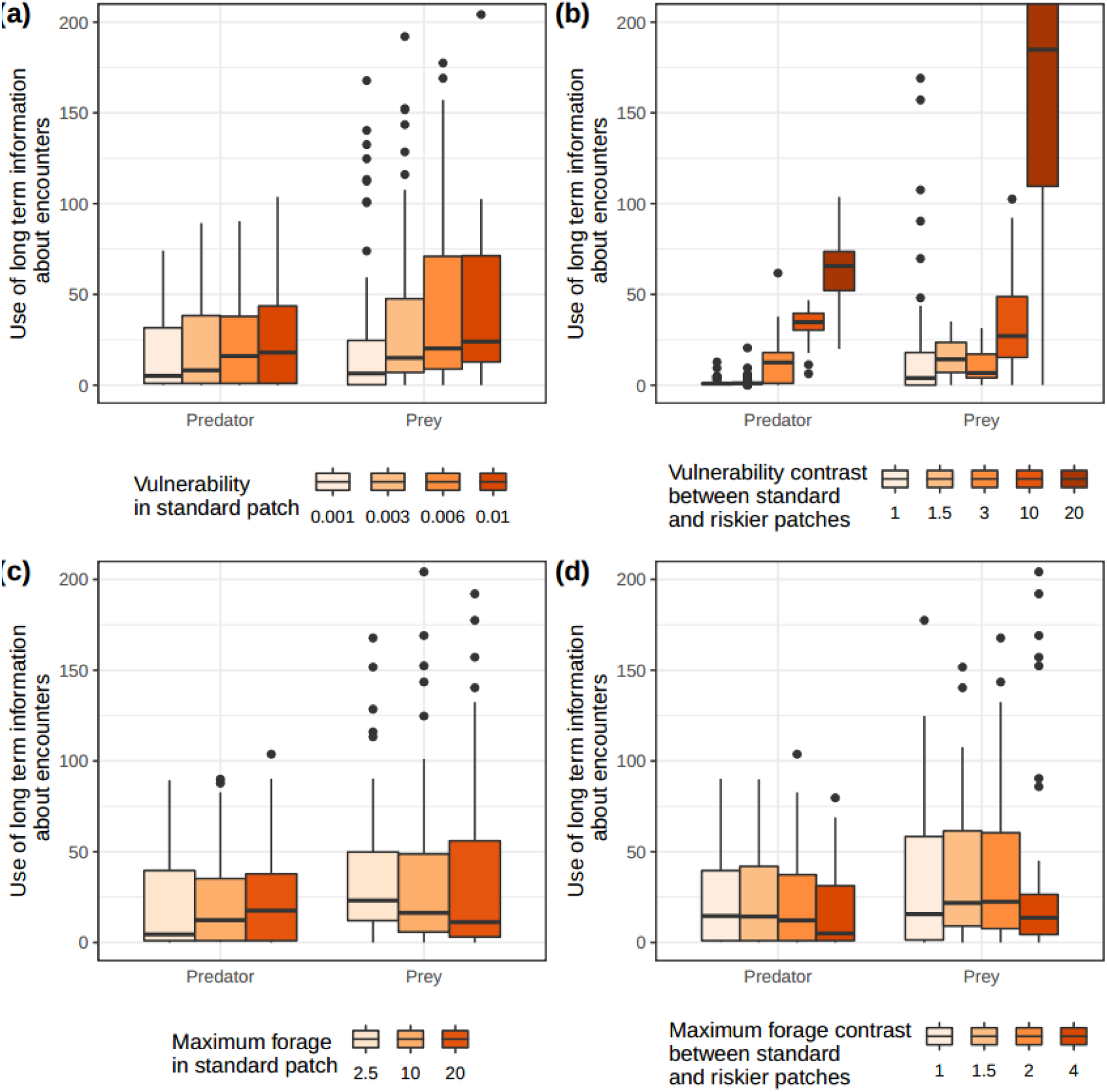
Optimal values, for predators and prey, of parameter δ, indexing the use of long-term information about encounters, for varying (a) vulnerability in the standard patches, (b) vulnerability contrast between standard and rich patches, (c) maximum forage in the standard patches and (d) maximum forage contrast between standard and rich patches. Lower values correspond to a lower weight given to older information.

### Relative importance of recent vs. older information about forage availability

Despite the large variability observed in the results, prey appear to use recent information on forage availability more than predators (Fig. 4, Fig. S2-16 to S2-20 in SI 2). Note that predators however rely less on information about forage availability than prey to determine where to move (see above). For predators, the importance of recent vs older information on forage availability increases as vulnerability in standard patches decreases (Fig. 4a, Fig. S2-16), when vulnerability differs greatly between standard and riskier patches (Fig. 4b, Fig. S2-15), and as forage contrast between standard and riskier patches increases (Fig. 4d, Fig. S2-19). For prey, the importance of recent vs older information on forage availability increases as vulnerability in standard patches decreases (Fig. 4a, Fig. S2-16) and the vulnerability contrast between standard and riskier patches decreases (Fig. 4b, Fig. S2-17). Maximum forage in standard patches (Fig. 4c, Fig. S2-18) and contrast in maximum forage between standard and riskier patches (fig. 4d, Fig. S2-19) also affect how recent information is used by prey, but results varied largely between simulations. It is clear, however, that the absence of predators leads prey to mostly value recent information (Fig. 4).

**Figure 4.**
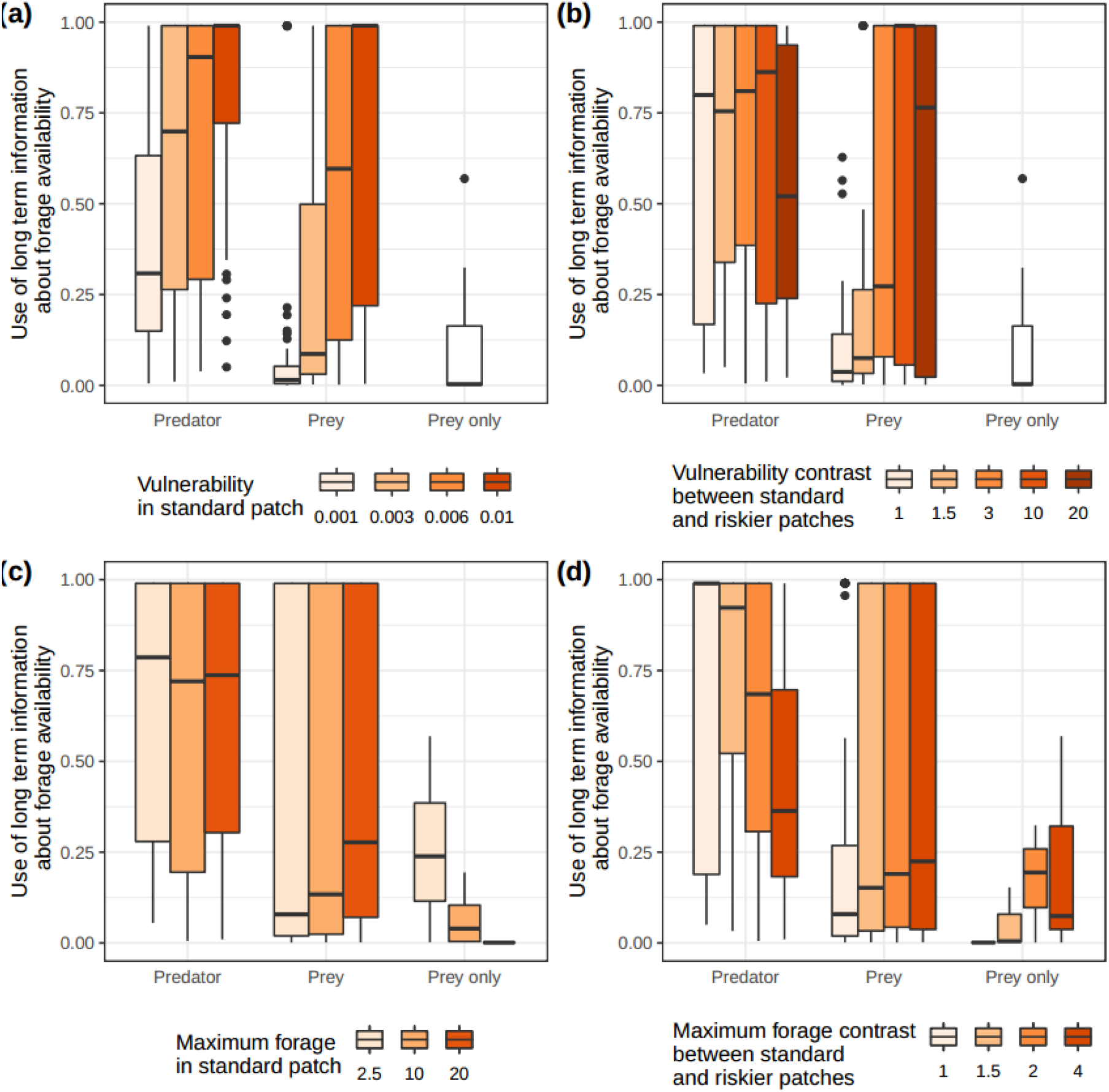
Optimal values, for predator and prey, of parameter μ, indexing the use of long-term information about forage availability, for varying (a) vulnerability in the standard patches, (b) vulnerability contrast between standard and rich patches, (c) maximum forage in the standard patches and (d) maximum forage contrast between standard and rich patches. Values for environments without predators are also shown (patches were then not assigned vulnerability levels). Lower values correspond to a lower weight given to older information.

## Discussion

We developed, to our knowledge, the first model predicting how optimal predators and prey should use information about prey forage availability and encounters with their enemies. The model revealed general differences in the optimal use of information between predators and prey, as well as the influence on patch choice strategies of key environmental characteristics such as contrasts in forage availability and vulnerability between patches. Our study offers a theoretical framework to interpret previous studies and generalize their predictions. In particular, our model provides three clear predictions that apply to the realistic situation of competitive prey foraging under predation risk: (1) predators and prey play a shell game in which each player is trying to keep its location unpredictable. Predators remain however much more unpredictable than prey; (2) what makes a good patch for a predator is more linked to successful encounters with prey than high prey forage availability; the opposite is true for prey; (3) prey and predators used recent and old information about encounters rather similarly, and give more weight to recent encounters. In contrast, the relative importance of recent and old information about forage availability is highly variable, differing between prey and predators and depending on environmental characteristics. We now discuss these predictions further, in the light of existing and possible studies.

Previous theoretical (Mitchell & Lima 2002; Mitchell 2009) and empirical (Roth & Lima 2007; Valeix et al. 2011; Bosiger et al. 2012; Simon et al. 2019) studies have suggested that predators and prey play a shell game in which both the prey and the predator would benefit from moving regularly in an unpredictable manner to prevent the other player to learn and then avoid (for the prey) or focus on (for the predator) places where the other player would be. Theoretical studies have however focused on unpredictability of the prey, as they never allowed predators to move somewhat randomly. Empirical studies never compared movement predictability of predators and prey. Our model fills these gaps from a theoretical standpoint, and predicts that predators’ movements should often be unpredictable. A prey knowing a predator’s patch weights would not be able to predict its movement as the predator’s movements were only weakly determined by those weights. Conversely, although prey did not always move towards the best patches, a predator knowing prey patch weights would significantly increase its chances to find prey, as it could restrict the number of patches where to look for, as prey most often selected patches with a probability at least proportional to the patch weight. The comparison of prey strategies in presence or absence of predators however clearly demonstrates that, in absence of predators, prey would be much more predictable, always moving towards the patch perceived as best. The general contrast in movement randomness between predators and prey is robust to variations in the levels and contrasts of forage availability and vulnerability in the landscape.

Differences in movement strategies between predators and prey could be explained by considering the costs and benefits of randomness for both. Regarding benefits, random movement by predators increases the availability of prey that, if behaviourally responsive to predictable risk, could otherwise avoid places where predators would be likely to be present, leading to behavioural resource depression (sensu Charnov 1976). Random movement by prey benefits them as it reduces the searching efficiency of predators that can memorize past encounters and could use this information for later searches (Mitchell 2009). As for costs, even though predators could recognize a good patch, mostly based on past encounters as high forage availability is just a proxy for the potential presence of prey (Flaxman & Lou 2009), the cost of ignoring this information and moving randomly is likely to be lower than for the prey. Predators moving randomly and arriving at a bad patch only suffer from missed opportunity costs of foraging. In contrast, prey moving randomly rather than doing an informed choice and arriving on a bad patch could suffer from two costs, lower forage intake and increased predation risk. In our simulations, prey movement was virtually always strongly determined by patch weights, suggesting that in most circumstances the costs associated with randomness largely outweighed its potential benefits. In addition, as predators evolve to be unpredictable, information about past encounters becomes less and less relevant to the prey. Prey therefore end up giving much more weight to information about forage availability than to encounters, focusing their movement towards good foraging patches, with a low, but not negligible, level of unpredictability to extend the predator’ area to search, giving prey a chance to avoid encounters. Overall, with our model, we confirm the potential importance of randomness in the predator-prey spatial game, and explain why we should expect predators to be unpredictable, and prey not so, but more than in absence of predation.

Our model also provides insights on how prey and predators should use recent and past information about encounters and forage availability. Recent encounters appear to be given more weight than older ones by both prey and predators. This make sense as, generally, information about encounters is poorly used by both: prey because they cannot predict where predators occur and thus mostly use information about forage availability, predators because they move rather randomly and therefore do not use much information, although when they do they favor information about where they met prey rather than about prey forage availability. It is likely that prey and predators used information about recent encounters to escape (prey) or remain in (predator) patches where an encounter occurred. It is consistent with observations that prey often leave patches when a predator is detected, and do not return immediately (Courbin et al. 2016). Predators are likely to leave rapidly anyway as patch weight has a low contribution to patch choice. In contrast, and despite strong variability among simulations and environmental contexts, recent and older information about forage availability was used differently by prey and predators. Prey generally gave more weight to recent information about forage than predators did. This could be explained by the fact that recent information is important for the prey as it allows it to keep foraging in good foraging patches, or move away when foraging in poor patches. For predators, recent forage availability might be a poor predictor of finding prey that could have arrived there by chance, whereas longer-term information about forage availability in patches might be a predictor of the likelihood of prey’s presence or density. As discussed before, however, information about forage availability remains weakly used by predators, unless patches do not differ in vulnerability, in which case predators move less randomly and give slightly more weight to information about forage than usual (Fig. 1 and 2). Thus, although our study suggests that leap-frogging, i.e. when predators track prey resources rather than prey themselves (Sih 1998; Flaxman & Lou 2009), should be rare, it might be more detectable in environments where prey vulnerability is homogeneous, as in the experimental contexts where it has been tested (Hammond et al. 2007; Williams & Flaxman 2012).

Generally, although the level and distribution of risk and prey forage in the environment did not qualitatively alter the observed differences between predators and prey space use strategies, it still had noticeable effects. Increasing global risk levels, by increasing the prey vulnerability in all patches, or only in riskier patches, leads prey to give increasing importance to encounters relative to forage availability. As risk increases, prey initially become slightly less predictable, but this effect does not persist, and possibly reverse, in very risky environments. Simultaneously, prey spend increasingly more time in standard, safer, patches (Fig. S3 in SI 3), increasingly using long-term information on encounter risk to identify these. Predators become less predictable and use patches increasingly randomly, although this effect appears to be slightly reduced when the vulnerability contrast between standard and riskier patches increases, predators then using long-term information about encounters, likely to avoid random movements that would take them to riskiest patches barely ever used by prey. Therefore, our results suggest that lowest levels of movement predictability of prey and predators should be observed at intermediate levels of vulnerability heterogeneity within the landscape. To the best of our knowledge, this theoretical prediction has not been made before, and is amenable to testing. Effects of the levels and heterogeneity of prey forage availability differed from those observed for vulnerability, and in cases affected movement predictability of prey and predators in a contrasted way. While predators become slightly less predictable as prey forage availability increases in all patches, prey become increasingly predictable, focusing more on standard, safest, patches, using long-term information on forage to identify them. Note how, in absence of predation, prey would in contrast increasingly use the most recent information to find most profitable patches. Prey predictability however decreases as riskier patches become more profitable. This occurs because the costs of not moving to the best patches is increasingly reduced by finding higher forage availability when the prey move to a riskier patch, which they increasing do (Fig. S3).

Overall, our work provides the most comprehensive theoretical framework to date to predict how prey and predator movement strategies should have evolved in contrasted environments. In particular, we focused on improving previous approaches by considering both prey and predators as behaviorally-responsive players of the space race, as this is now well established empirically (e.g. invertebrates: Hammond et al. 2007; large mammals: Valeix et al. 2011; Courbin et al. 2019). Our predictions should now be challenged by experimental and observational studies to reveal the extent of our understanding of predator-prey interactions. A key insight gained here is that the interplay between memory use and purposeful randomness should be a central focus of future studies.

## Acknowledgments

This work was supported by grant ANR-16-CE02-0001 from the ‘Agence Nationale de la Recherche’ to S. Chamaillé-Jammes, a grant from the Natural Science and Engineering Research Council to D. Fortin, and by the Montpellier Bioinformatics Biodiversity platform which provided computing time.

