## Supplementary material for "A theory of the use of information by enemies in the predator-prey space race": SI 1

### Supplementary Information 1

#### 1. Model parameters

Table S1. Symbol, definition and values of model parameters.

| Symbol | Definition | Equation | Value or range |
|---|---|---|---|
| **Parameters defining movement strategies (whose values are left to evolve towards optimal values using the genetic algorithm)** | | | |
| $\mu$ | Relative contribution of recent vs. older information about prey forage availability in determining patch weights. Lower values correspond to a lower importance given to older information | 5 | 0.001 – 0.99 |
| $\beta$ | Relative importance of encounters vs. prey forage availability in determining patch weights. Lower values correspond to lower importance given to encounters. | 6.1, 6.2 | 0.001 – 100 |
| $\delta$ | Relative contribution of recent vs. older information about encounters in determining patch weights. Lower values correspond to a lower importance given to older information. | 6.2 | 0 – 1e5 |
| $\alpha$ | Contribution of patch weights to the choice of the patches used. Lower values correspond to less informed, i.e. more random, movements. | 8 | 0.001 – 8 |
| **Parameters used to contrast patches** | | | |
| $K_i$ | Maximum forage (forage carrying capacity) in patch *i* | 3 | 2.5, 10, 20 for standard patches; values for riskier patches were 1, 1.5, 2 or 4 times greater |
| $\gamma_i$ | Prey vulnerability in patch *i*. The maximum value of vulnerability given in an environment, i.e. the vulnerability in riskier patches, is $\gamma_{max}$. | 6.1 | 0.001, 0.003, 0.006, 0.01 for standard patches; values for riskier patches were 1, 1.5, 10, 3, 10 or 20 times greater |

| Other parameters | | | |
|---|---|---|---|
| $r_m$ | Maximum growth rate of forage in patches | 3 | 0.2 |
| $R_{min}$ | Minimum forage in a patch at the beginning of a time-step | 3 | 0.1 |
| a | Prey attack rate of forage | 4.1 | 0.08 |
| h | Prey handling time of forage | 4.1 | 0.75 |
| $p_e$ | Probability of encounter between a predator and a prey when in the same patch | substep 2 | 0.1 |

**2. Model initialization**

Each simulation is initialized as follows: at $t_0$, the beginning of each simulation, 5 predators and 20 prey are randomly distributed among 20 patches (15 of them being standard patches). For each patch *i*, the amount of forage is set to its maximum value $K_i$. We assumed that predators and prey know all patches, and weights $W_i$ are initialized with identical values for all patches, using the following values for $F_i$ and $E_i$: for prey, $F_i$ are set to the mean forage intake calculated over all patches at $t_0$; for predators, $F_i$ is set to the mean forage availability calculated over all patches at $t_0$. $E_i$ are set to 1 for all predators and prey.
