## Supplementary material for "A theory of the use of information by enemies in the predator-prey space race": SI 2

### Supporting Information 2

This document provides information to evaluate the convergence of the genetic algorithm. Figures present the final values obtained by the genetic algorithm procedure for the parameters defining movement strategies, for each of the 3 replicates (termed chains). Results from chain 1 are those presented in the main text.

#### 1.1 Movement Predictability (parameter α)

Figure S2.1: Optimal values, for predators and prey, of parameter α, indexing movement predictability, for varying vulnerability in the standard patches. Values for environments without predators are also shown (patches were then not assigned vulnerability levels). Lower values correspond to less informed, i.e. more random, movements.

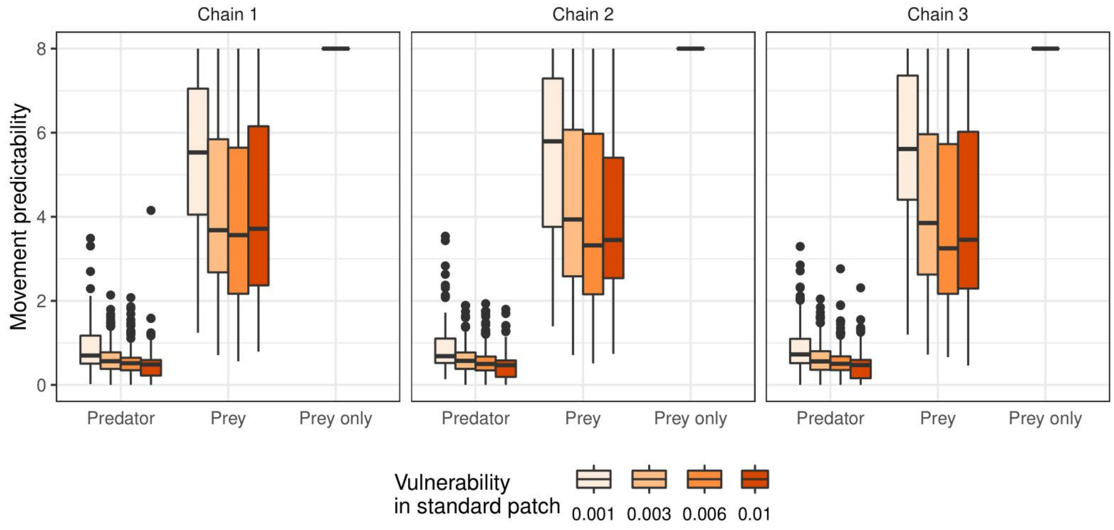


Figure S2.2: Optimal values, for predators and prey, of parameter α, indexing movement predictability, for varying vulnerability contrast between standard and rich patches. Values for environments without predators are also shown (patches were then not assigned vulnerability levels). Lower values correspond to less informed, i.e. more random, movements.

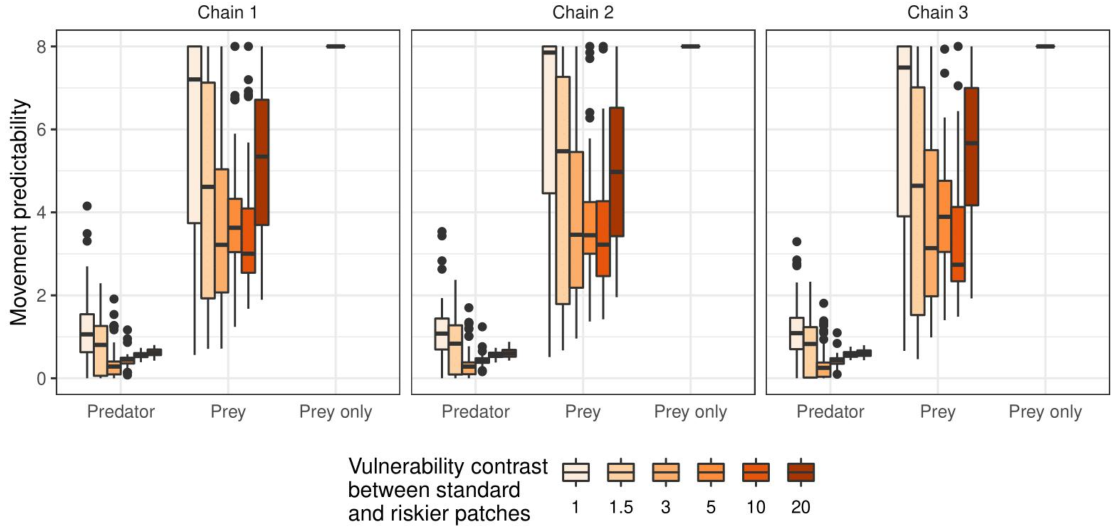

Figure S2.3: Optimal values, for predators and prey, of parameter α, indexing movement predictability, for varying maximum forage in the standard patches. Values for environments without predators are also shown (patches were then not assigned vulnerability levels). Lower values correspond to less informed, i.e. more random, movements.

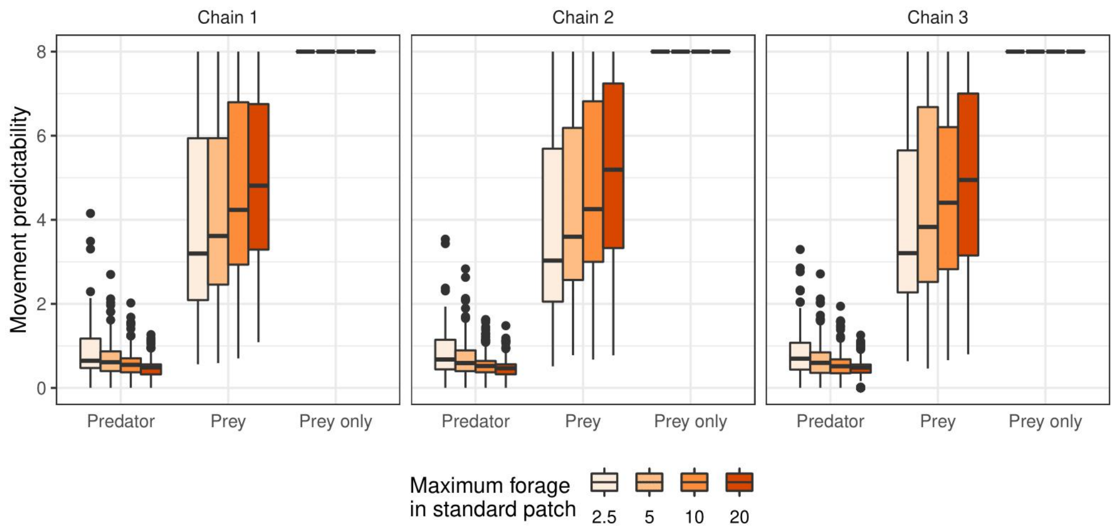


Figure S2.4: Optimal values, for predators and prey, of parameter α, indexing movement predictability, for varying maximum forage contrast between standard and rich patches. Values for environments without predators are also shown (patches were then not assigned vulnerability levels). Lower values correspond to less informed, i.e. more random, movements.

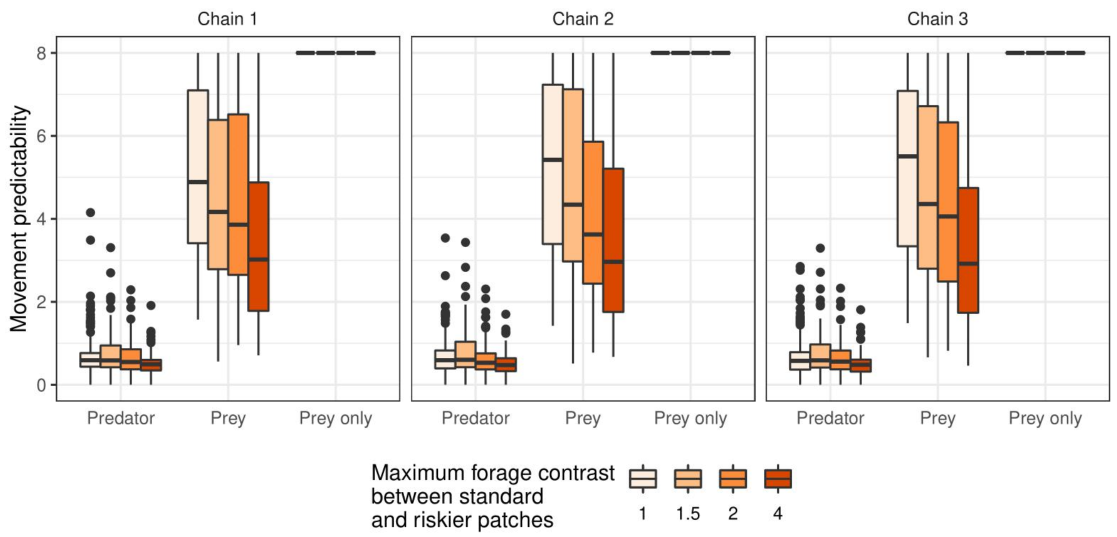

Figure S2.5: Optimal values of parameter $\alpha$, indexing movement predictability, for the 240 different environments tested and the 3 replicates (chains). Each small square represents a different environment type characterized by the maximum forage in the standard patches ($K_i$), the maximum forage contrast between standard and richer patch (Contrast($K_i$)), the vulnerability in the standard patch ($\gamma_i$), and the vulnerability contrast between standard and riskier patch (Contrast($\gamma_i$)). Values are shown for both predators (upper triangle) and prey (lower triangle). Lower values correspond to less informed, i.e. more random, movements.

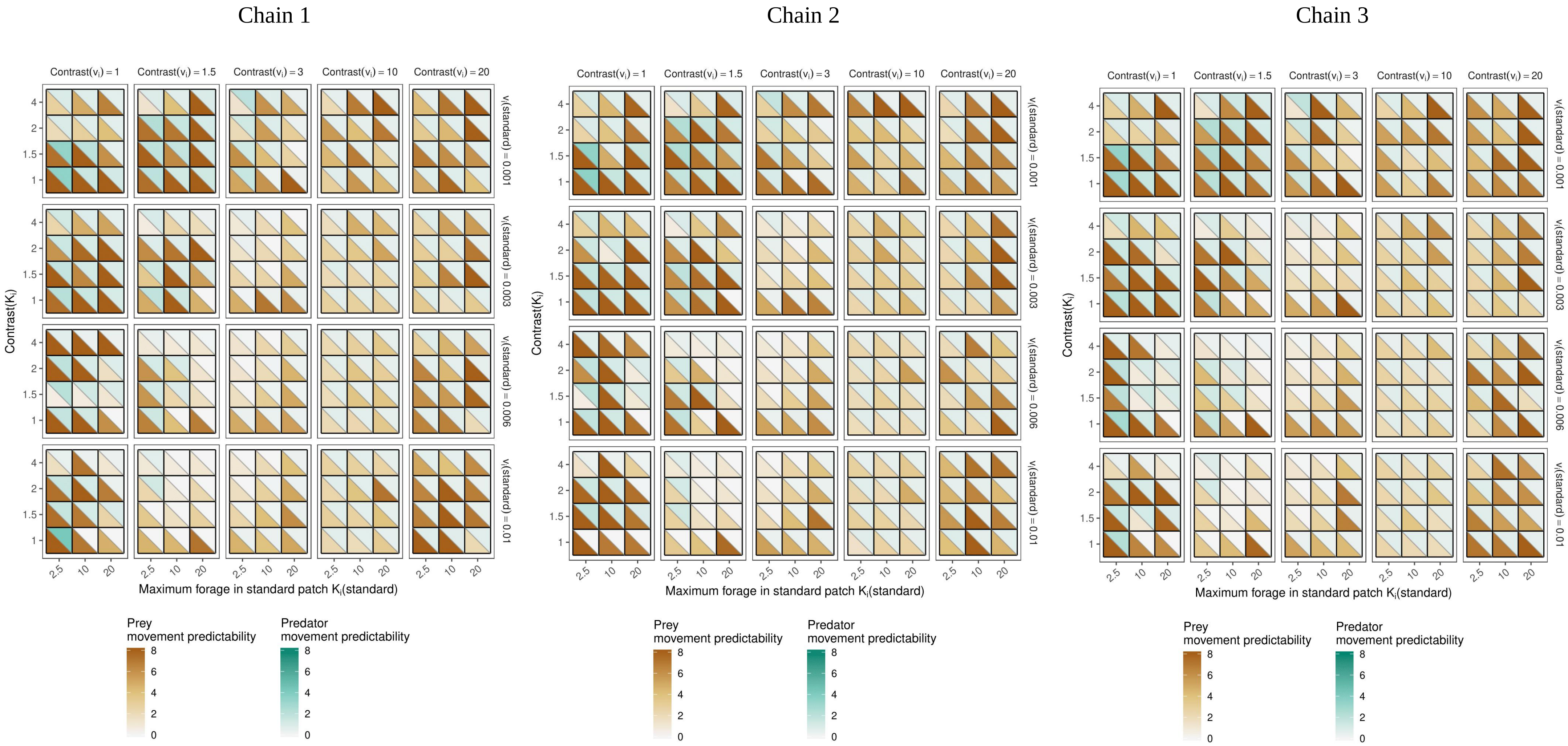

#### 1.2 Relative importance of information about encounters and forage availability (β)

Figure S2.6: Optimal values, for predators and prey, of parameter β, indexing the relative importance of information about encounters, for varying vulnerability in the standard patches, (b) vulnerability contrast between standard and rich patches, (c) maximum forage in the standard patches and (d) maximum forage contrast between standard and rich patches. Lower values correspond to lower importance given to encounters, relative to prey forage availability.

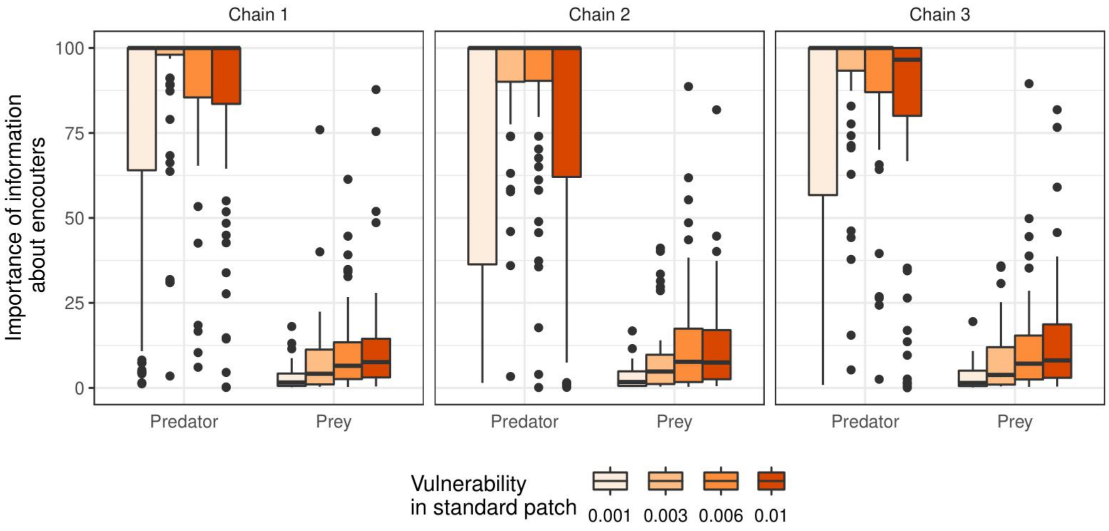


Figure S2.7: Optimal values, for predators and prey, of parameter β, indexing the relative importance of information about encounters, for varying vulnerability contrast between standard and rich patches. Lower values correspond to lower importance given to encounters, relative to prey forage availability.

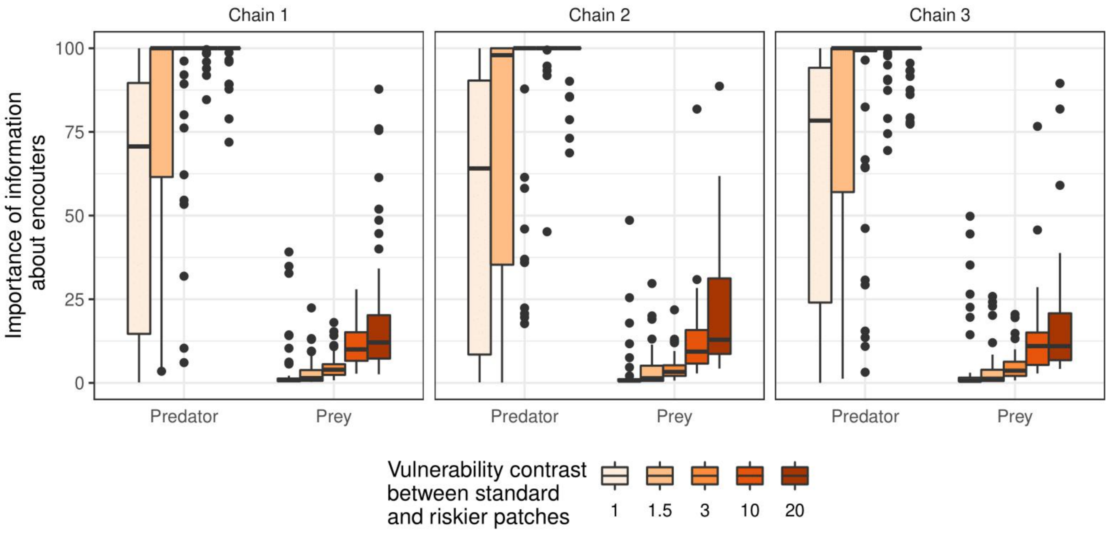

Figure S2.8: Optimal values, for predators and prey, of parameter β, indexing the relative importance of information about encounters, for varying maximum forage in the standard patches. Lower values correspond to lower importance given to encounters, relative to prey forage availability.

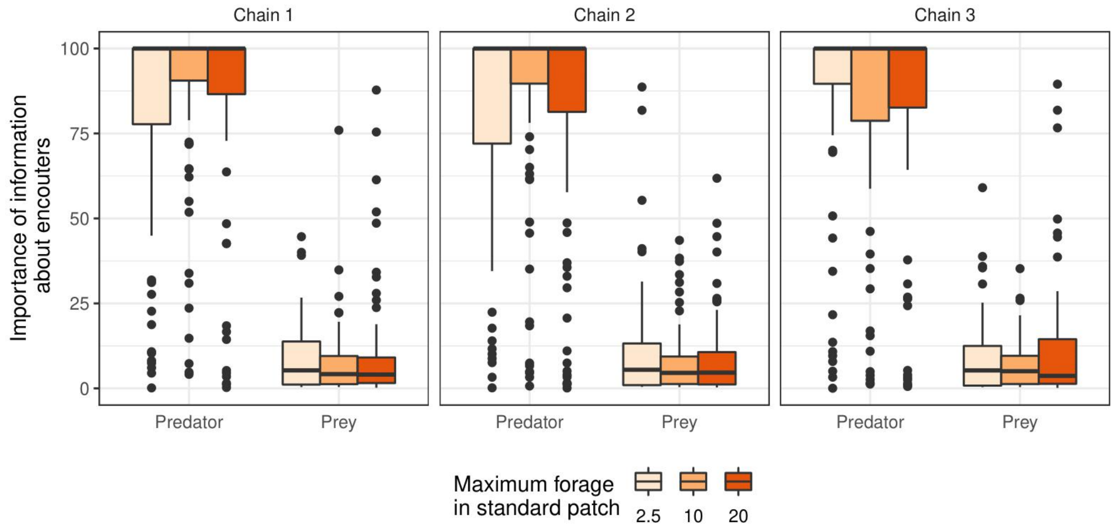


Figure S2.9: Optimal values, for predators and prey, of parameter β, indexing the relative importance of information about encounters, for varying maximum forage contrast between standard and rich patches. Lower values correspond to lower importance given to encounters, relative to prey forage availability.

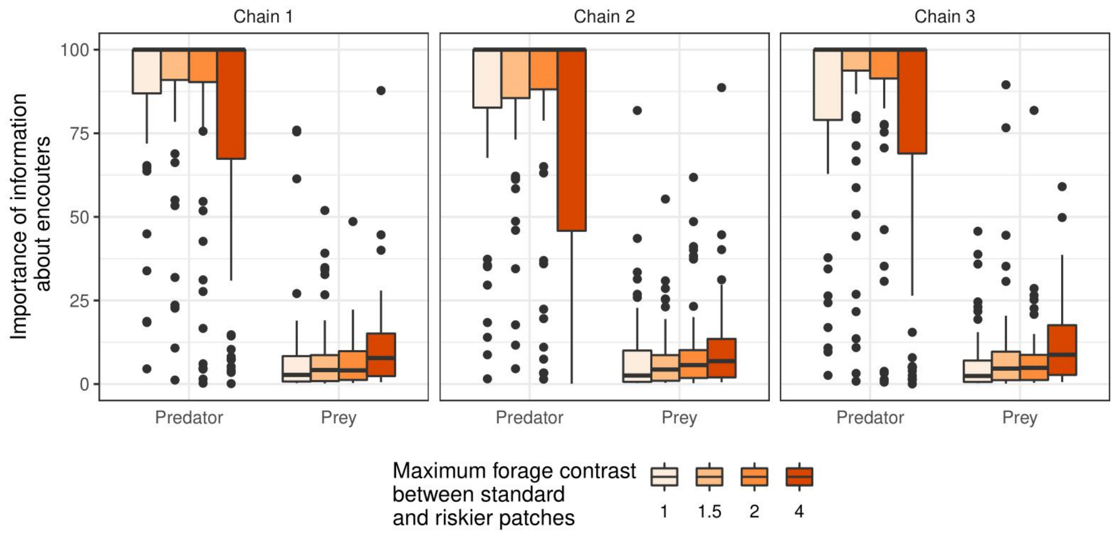

Figure S2.10: Optimal values of parameter β, indexing the relative importance of information about encounters, for the 240 different environments tested and the 3 replicates (chains). Each small square represents a different environment type characterized by the maximum forage in the standard patches ($K_i$), the maximum forage contrast between standard and richer patch (Contrast($K_i$)), the vulnerability in the standard patch ($\gamma_i$), and the vulnerability contrast between standard and riskier patch (Contrast($\gamma_i$)). Values are shown for both predators (upper triangle) and prey (lower triangle). Lower values correspond to lower importance given to encounters, relative to prey forage availability.

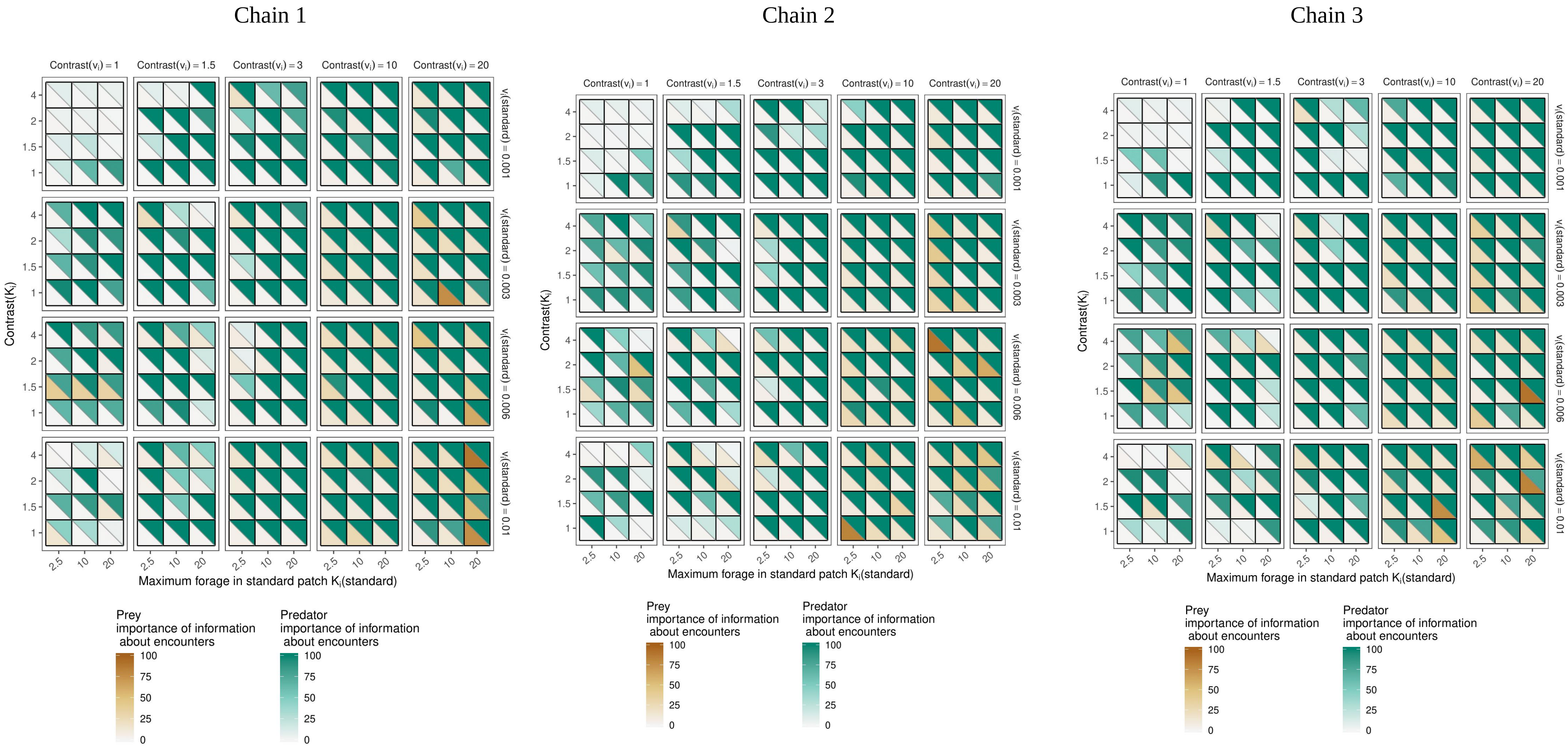

#### 1.3 Use of long term information about encounters (δ)

Figure S2.11: Optimal values, for predators and prey, of parameter δ, indexing the use of long-term information about encounters, for varying vulnerability in the standard patches. Lower values correspond to a lower weight given to older information.

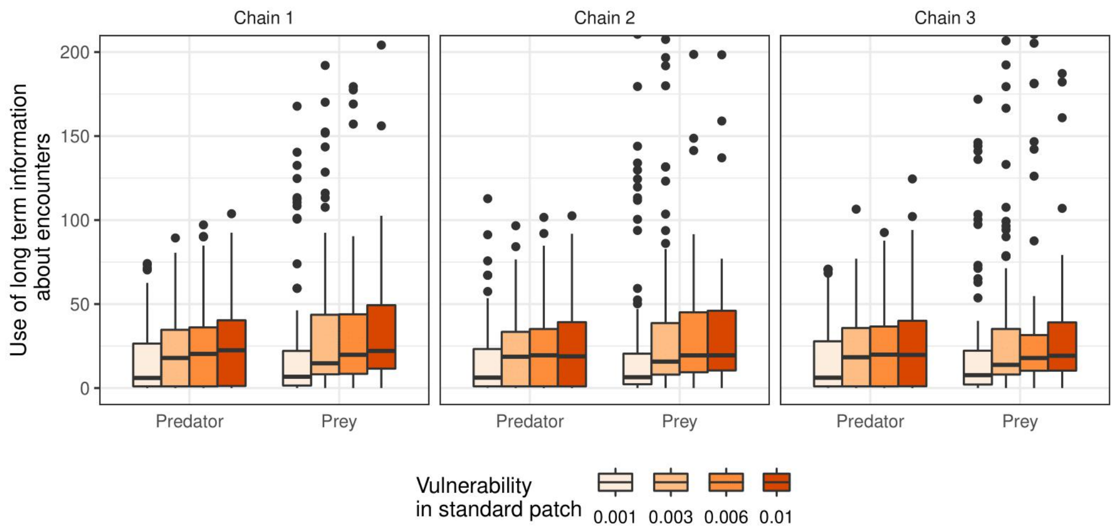


Figure S2.12: Optimal values, for predators and prey, of parameter δ, indexing the use of long-term information about encounters, for varying vulnerability contrast between standard and rich patches. Lower values correspond to a lower weight given to older information.

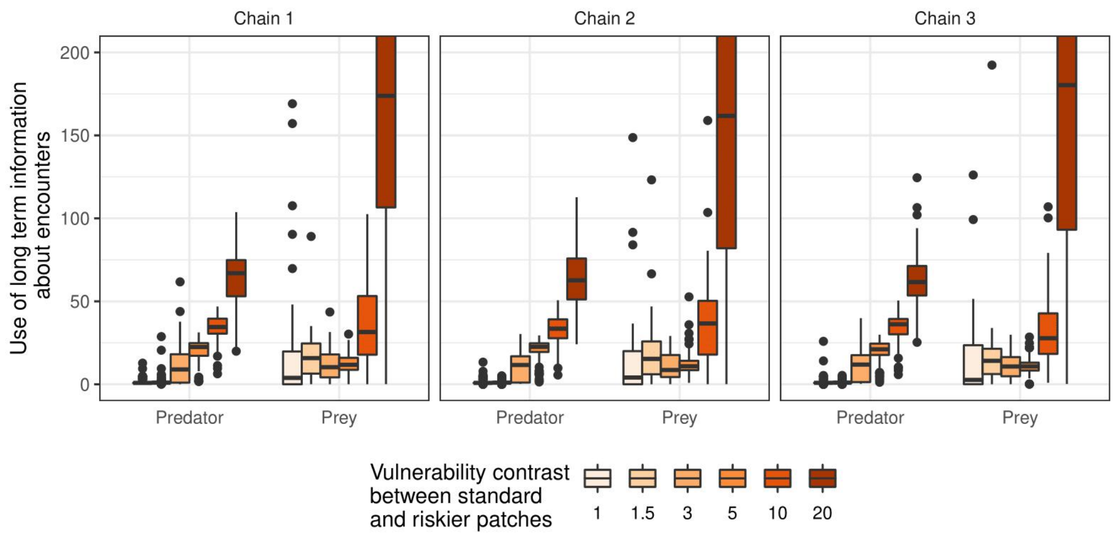

Figure S2.13: Optimal values, for predators and prey, of parameter δ, indexing the use of long-term information about encounters, for varying maximum forage in the standard patches. Lower values correspond to a lower weight given to older information.

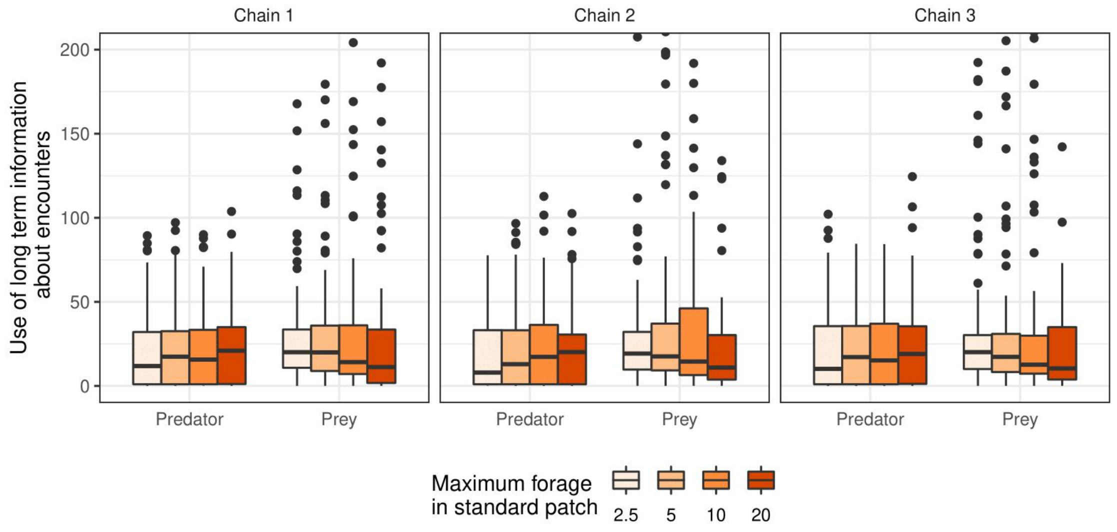


Figure S2.14: Optimal values, for predators and prey, of parameter δ, indexing the use of long-term information about encounters, for varying maximum forage contrast between standard and rich patches. Lower values correspond to a lower weight given to older information.

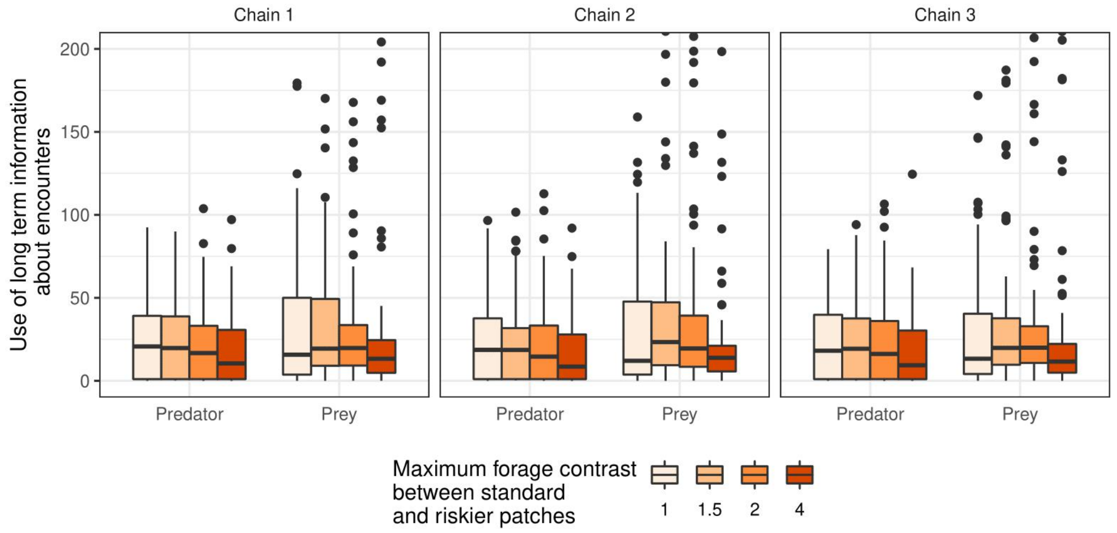

Figure S2.15: Optimal values of parameter δ, indexing the use of long-term information about encounters, for the 240 different environments tested and the 3 replicates (chains). Each small square represents a different environment type characterized by the maximum forage in the standard patches ($K_i$), the maximum forage contrast between standard and richer patch (Contrast($K_i$)), the vulnerability in the standard patch ($\gamma_i$), and the vulnerability contrast between standard and riskier patch (Contrast($\gamma_i$)). Values are shown for both predators (upper triangle) and prey (lower triangle). Lower values correspond to a lower weight given to older information.

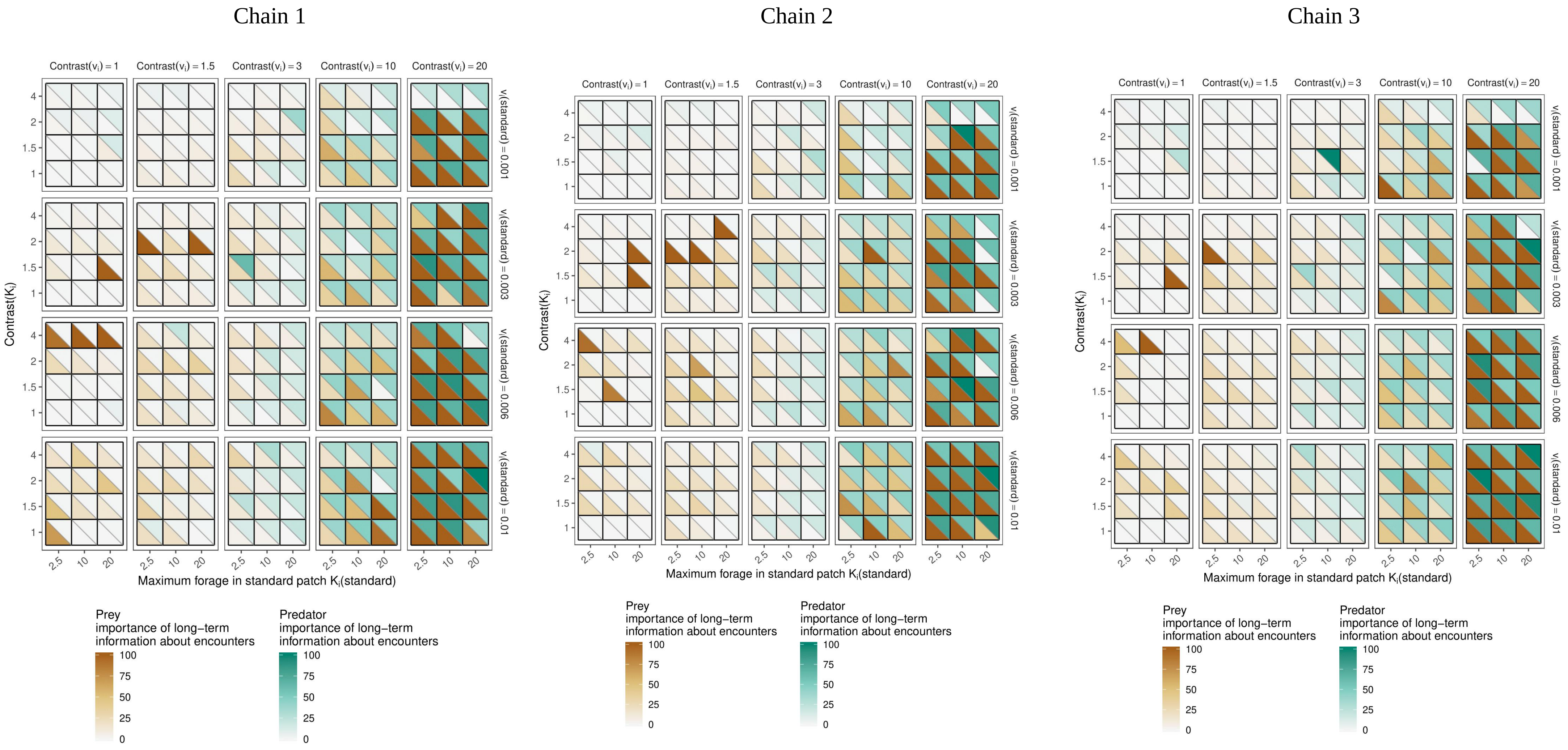

#### 1.4 Use of long term information about forage availability (μ)

Figure S2.16: Optimal values, for predator and prey, of parameter μ, indexing the use of long-term information about forage availability, for varying vulnerability in the standard patches. Values for environments without predators are also shown (patches were then not assigned vulnerability levels). Lower values correspond to a lower weight given to older information.

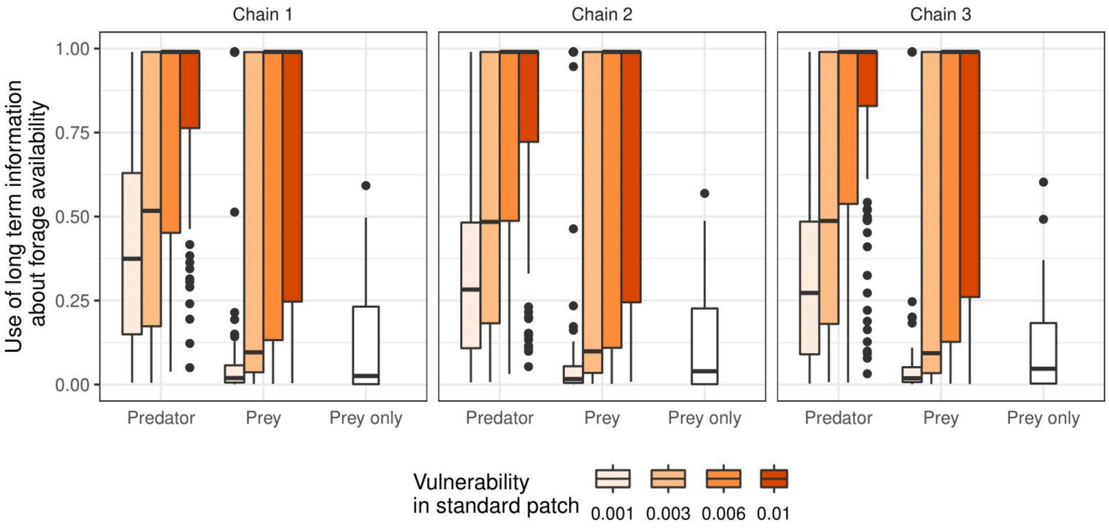


Figure S2.17: Optimal values, for predator and prey, of parameter μ, indexing the use of long-term information about forage availability, for varying vulnerability contrast between standard and rich patches. Values for environments without predators are also shown (patches were then not assigned vulnerability levels). Lower values correspond to a lower weight given to older information.

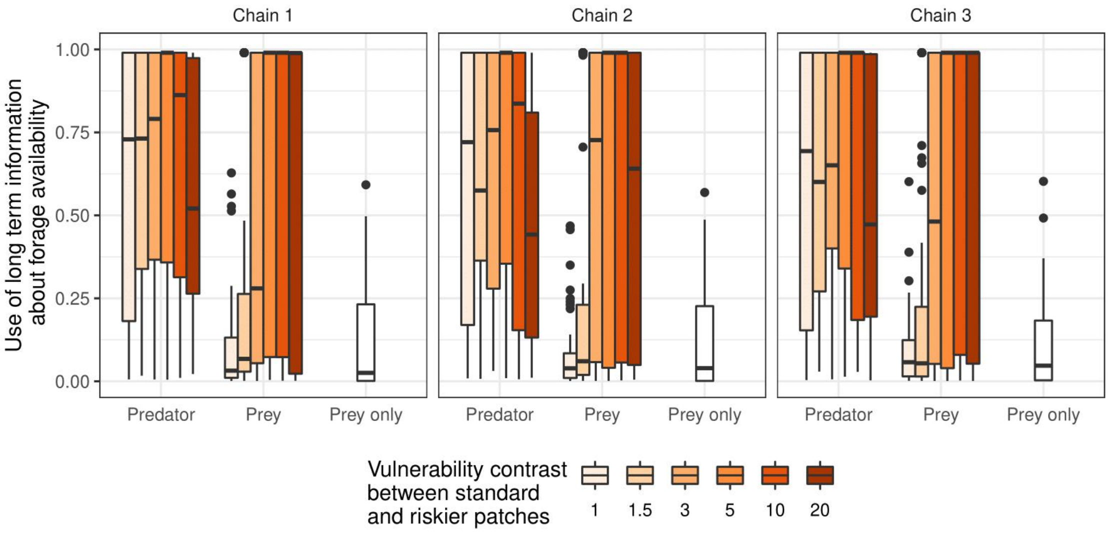

Figure S2.18: Optimal values, for predator and prey, of parameter μ, indexing the use of long-term information about forage availability, for varying maximum forage in the standard patches. Values for environments without predators are also shown (patches were then not assigned vulnerability levels). Lower values correspond to a lower weight given to older information.

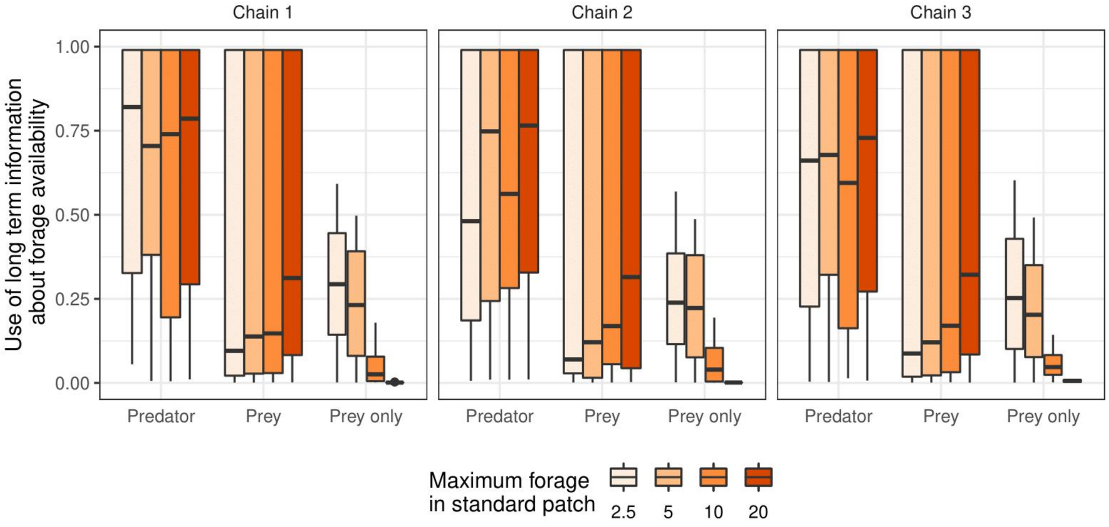


Figure S2.19: Optimal values, for predator and prey, of parameter μ, indexing the use of long-term information about forage availability, for varying maximum forage contrast between standard and rich patches. Values for environments without predators are also shown (patches were then not assigned vulnerability levels). Lower values correspond to a lower weight given to older information.

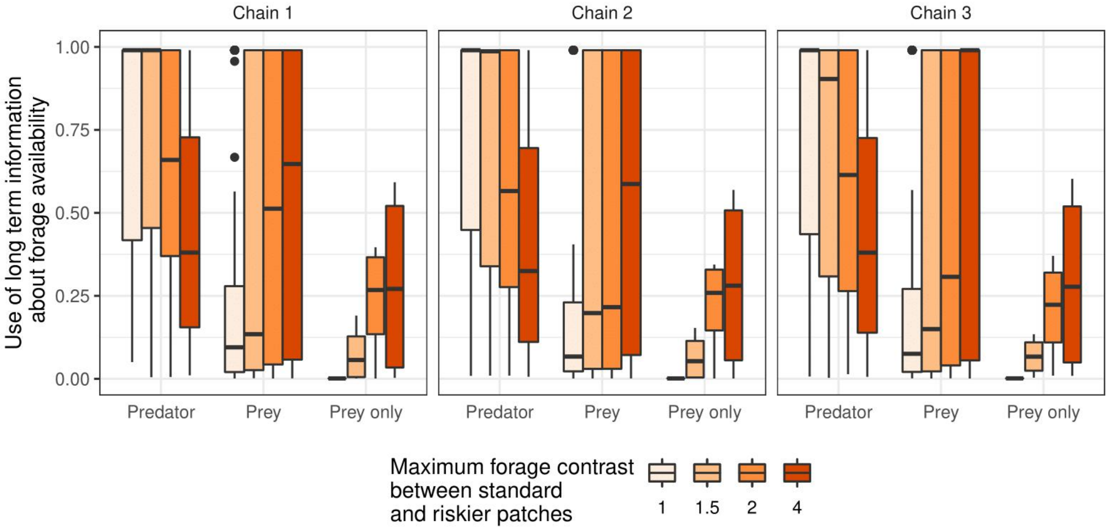

Figure S2.20: Optimal values of parameter μ, indexing the use of long-term information about forage availability, for the 240 different environments tested and the 3 replicates (chains). Each small square represents a different environment type characterized by the maximum forage in the standard patches ($K_i$), the maximum forage contrast between standard and richer patch (Contrast($K_i$)), the vulnerability in the standard patch ($\gamma_i$), and the vulnerability contrast between standard and riskier patch (Contrast($\gamma_i$)). Values are shown for both predators (upper triangle) and prey (lower triangle). Lower values correspond to a lower weight given to older information.

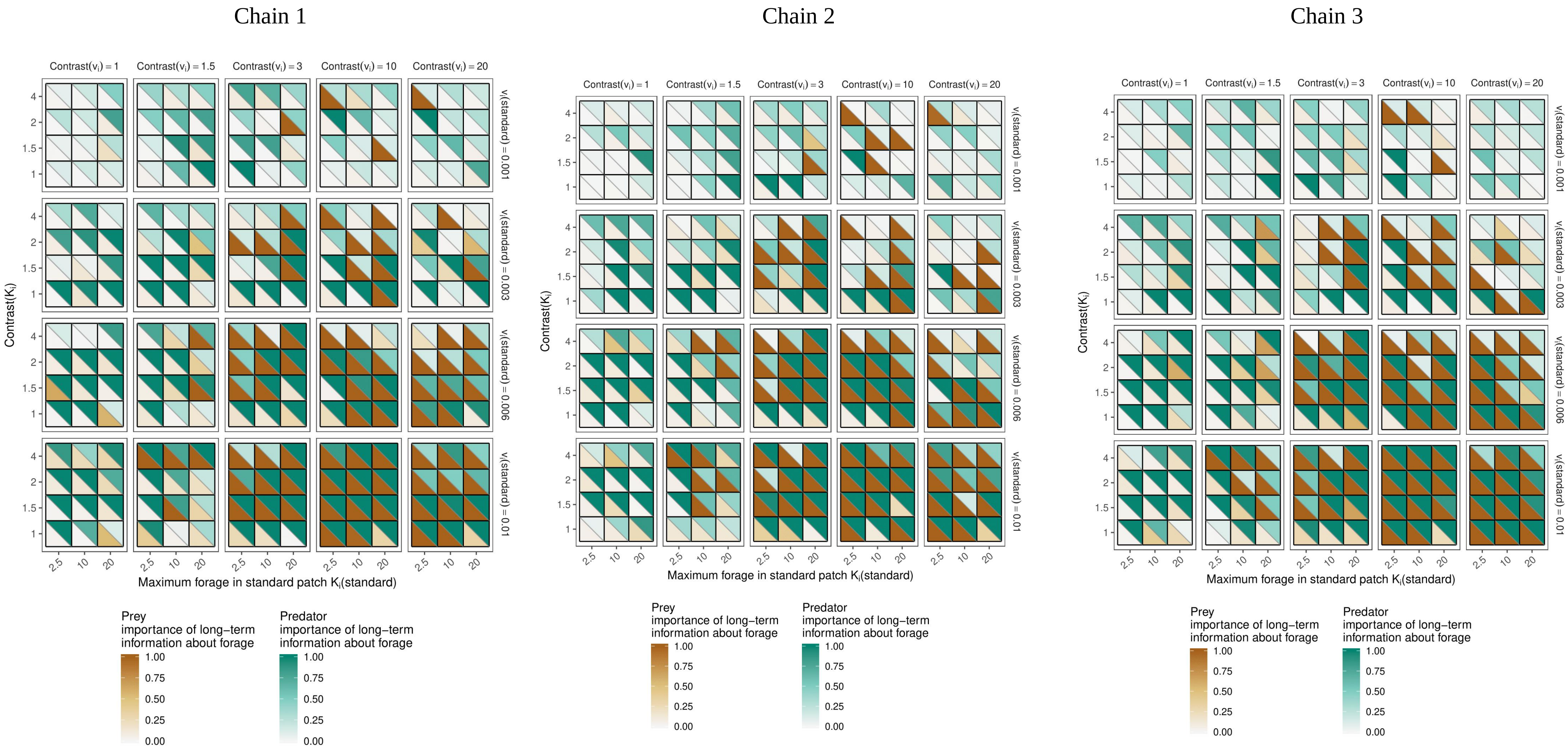
