## Supplementary material for "A theory of the use of information by enemies in the predator-prey space race": SI 3

### Supporting Information 3

Figure S3. Probability of presence in a risky patch, for both predators and prey, for the 240 different environments tested and the 3 replicates (chains). Each small square represents a different environment type characterized by the maximum forage in the standard patches ($K_i$), the maximum forage contrast between standard and richer patch (Contrast($K_i$)), the vulnerability in the standard patch ($\gamma_i$), and the vulnerability contrast between standard and riskier patch (Contrast($\gamma_i$)). Within each small square, the color is linked to the probability that a predator (upper triangle) or a prey (lower triangle) is located at time t in a risky patch. White indicates homogeneous space use, with probability being 0.25 because 25% of the patches are risky ones (SI 1).

Chain 1

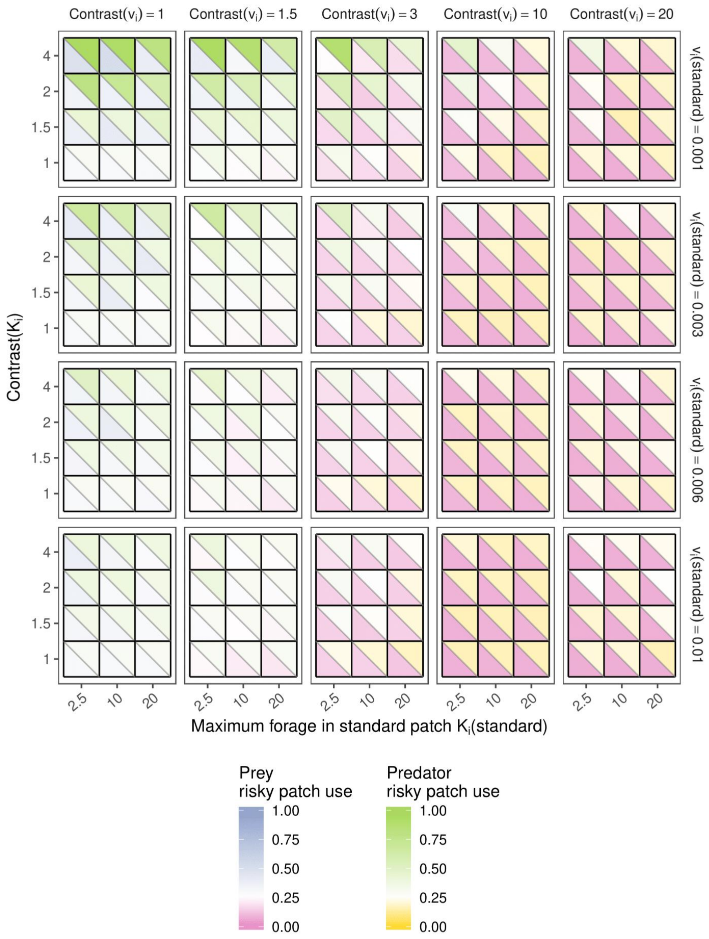


Chain 2

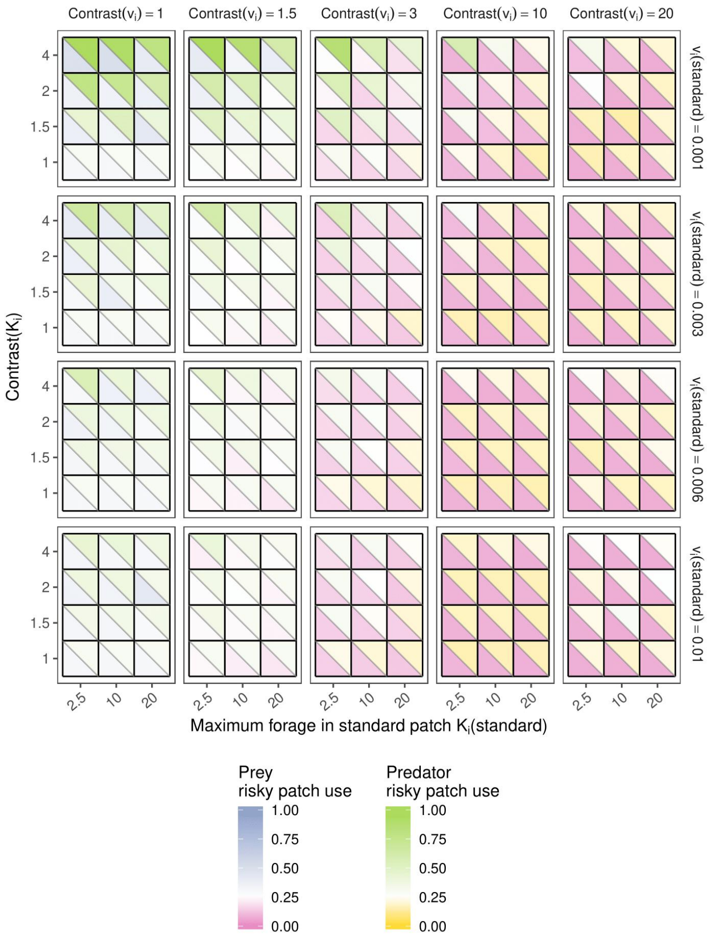


Chain 3

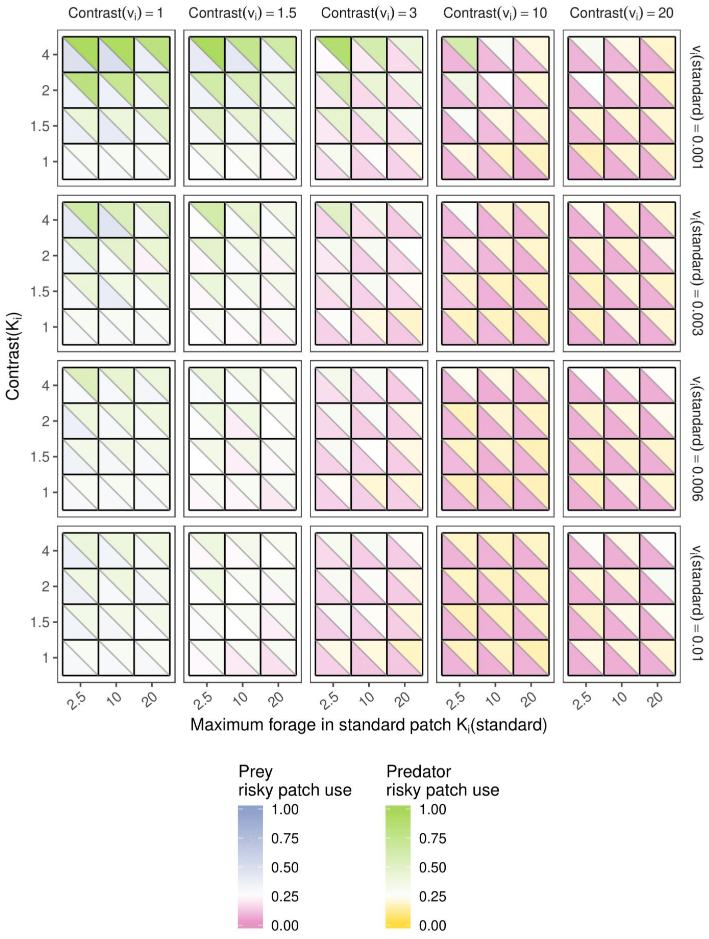
